# A single *Vibrionales* 16S rRNA oligotype dominates the intestinal microbiome in two geographically separated Atlantic cod populations

**DOI:** 10.1101/186346

**Authors:** Even Sannes Riiser, Thomas H.A. Haverkamp, Ørnulf Borgan, Kjetill S. Jakobsen, Sissel Jentoft, Bastiaan Star

## Abstract

**Background:** Host-microbe interactions are particularly intriguing in Atlantic cod (*Gadus morhua*), as it lacks the MHC II complex involved in presentation of extracellular pathogens. Nonetheless, little is known about the diversity of its microbiome in natural populations. Here, we use 16S rRNA high-throughput sequencing to investigate the microbial community composition in gut content and mucosa of 22 adult individuals from two coastal populations in Norway, located 470 km apart.

**Results:** We identify a core microbiome of 23 OTUs (97% sequence similarity) in all individuals that comprises 93% of the total number of reads. The most abundant orders are classified as *Vibrionales, Fusobacteriales, Clostridiales* and *Bacteroidales*. While mucosal samples show significantly lower diversity than gut content samples, no differences in OTU community composition are observed between the two populations. The differential abundance of oligotypes within two common OTUs does reveal limited spatial segregation. Remarkably, the most abundant OTU consists of a single oligotype (order *Vibrionales*, genus *Photobacterium*) that represents nearly 50% of the reads in both locations.

**Conclusions:** Our results show that the intestinal bacterial community of two geographically separated coastal populations of Atlantic cod is dominated by a limited number of highly abundant 16S rRNA oligotypes shared by all specimens examined. The ubiquity of these oligotypes suggests that the northern coastal Atlantic cod gut microbiome is colonized by a limited number of species with excellent dispersal capabilities that are well suited to thrive in their host environment.

## Background

Complex and specialized gut bacterial communities, collectively called the intestinal microbiome, provide a multitude of biological functions in fish (reviewed in [1–5]). For instance, the intestinal microbiome is essential in processes like fermentation in the hindgut of herbivorous species [6], innate immunity [7,8], vitamin synthesis [9] and influences expression of numerous host genes [10]. Despite these important biological roles, the diversity of the intestinal microbiome in natural populations remains unknown for many fish species. Studying microbial diversity in wild populations of teleosts provides valuable information regarding the factors affecting establishment and final composition of the fish gut microbiome [11]. This likely involves a complex interaction between exogenous factors (diet, salinity etc.) and endogenous factors (genotype, trophic level etc.) [3,12]. Comparative analyses of different fish species indicate that environmental variables (i.e. in salinity and trophic level) and fish taxonomy in particular contribute to the community composition in fish [4]. Intriguingly, in Atlantic salmon (*Salmo salar*) the bacterial community composition is not significantly different in populations sampled from both sides of the Atlantic ocean [13]. Whether such low spatial differentiation is the norm in other marine fish remains largely unexplored.

Atlantic cod (*Gadus morhua*) is an ecologically and commercially important species of the North Atlantic Ocean and represents a unique study system of the fish gut microbiome for several reasons. First, the gadiform adaptive immune system lacks the Major Histocompatibility Complex (MHC) II, which may affect its host-microbe interactions [14–16]. Second, this omnivore is exposed to a variety of environmental conditions due to its ability to exploit a wide range of ecological niches [17]. It has a large spatial distribution, which comprises various subpopulations with divergent migratory behavior [18]. Finally, the intestinal wall of Atlantic cod contains a large number of goblet cells producing prolific levels of mucosa, compared to Atlantic salmon [19]. The dense mucosal layer provides an opportunity for comparative analyses of microbial communities in the intestinal content vs. those of the mucosal layer [20].

Culture-based studies have demonstrated that the microbiome of Atlantic cod is dominated by *Vibrionales*, *Bacteroidales*, *Erysipelotrichales* and *Clostridiales* [21,22]. Nevertheless, high inter-individual variation in the intestinal bacterial community composition has been observed among reared cod-larvae [23]. Similarly, inter-individual variation has been observed at a single location, using culture-independent 16S rRNA sequencing [24]. These observations suggest that the final community composition is the result of a stochastic process during bacterial colonization, although little is known about the natural variation of the intestinal bacterial communities over larger geographical scales in Atlantic cod.

Our goal is to investigate the intestinal bacterial community composition in two coastal Atlantic cod populations located 470 kilometers apart using culture-independent high-throughput sequencing of the 16S rRNA V4 region. We compare the taxonomic composition and diversity of the Atlantic cod intestinal microbiome between two locations (Lofoten and Sørøya, Fig. 1a). Additionally, to ensure the classification of the intestinal bacterial community we investigate the microbiome composition of gut content vs. mucosa. We find that a single 16S rRNA *Vibrionales* oligotype (genus *Photobacterium*) comprises nearly 50% of the reads in both locations, and show that the intestinal bacterial community is numerically dominated by a limited amount of highly abundant 16S rRNA oligotypes. There are no significant differences in community composition between the two locations, while there is a small, but significant difference between gut content and mucosa. The ubiquity of the dominant 16S rRNA oligotypes suggests that the gut microbiome of the northern coastal Atlantic cod is colonized by a limited number of species with excellent dispersal capabilities that are well suited to thrive in their host environment.

**Figure 1:**
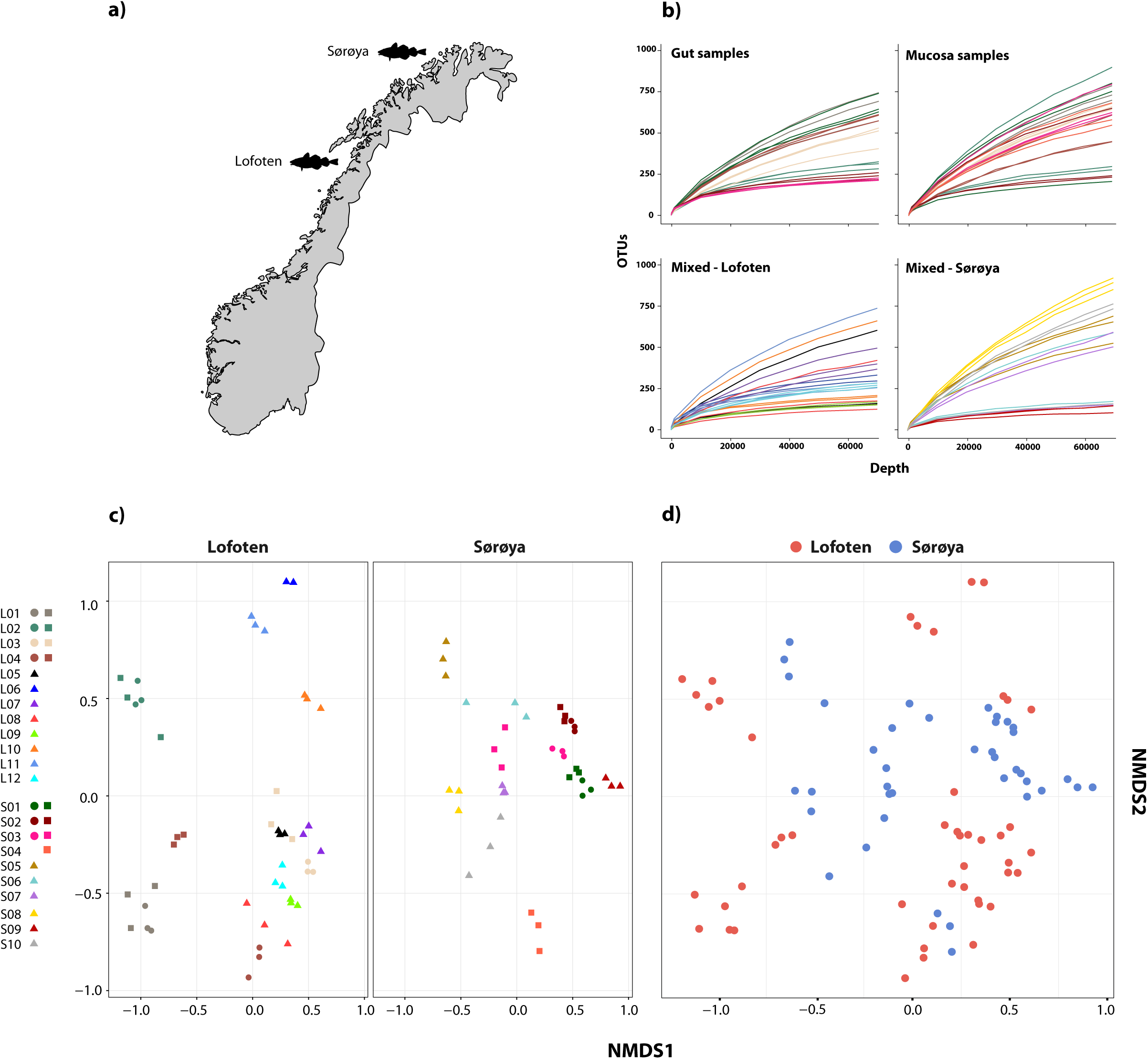
Microbial intestinal communities of wild Atlantic cod. **a) Map of sampling locations b)** Rarefaction curve analysis showing the number of detected OTUs as a function of read number (depth) for 22 Atlantic cod specimens. For seven specimens (upper two curve plots) gut and mucosal samples were successfully collected from the same individual and sequenced. Sequences are clustered using a 97% similarity cut-off. Replicates (3 per individual, except 2 for L06) are colored identically within each plot. Gut and mucosal samples collected from the same individual are colored identically. Color corresponds to the legend in c). All samples are normalized to the same maximum sequencing depth. **c)** Non-metric multidimensional scaling (NMDS) plot of all samples based on Bray-Curtis dissimilarity. Samples are plotted separately per location. Replicates from the same individual are colored identically. L = Lofoten, S = Sørøya. Gut (●), mucosa (■) and mixed (▲). **d)** Non-metric multidimensional scaling (NMDS) plot of Lofoten (red) and Sørøya (blue) based on Bray-Curtis dissimilarity.

## Methods

### Sample collection

Coastal Atlantic cod specimens were collected in Lofoten (N68.0619167, W13.5921667) (12 individuals, August 2014) and Sørøya (N70.760418, W21.782716) (10 individuals, September 2013) (Additional file 1). We obtained samples as a byproduct of conventional business practice, and all specimens were caught by commercial vessels, euthanized by local fishermen and were intended for human consumption. This sampling follows the guidelines set by the “Norwegian consensus platform for replacement, reduction and refinement of animal experiments” [25]. A 3 cm long part of the hindgut (immediately above the short, wider rectal chamber) was aseptically removed *post-mortem* by scalpel and stored on 70% ethanol. The samples were frozen (-20°C) for long-term storage. Otoliths were collected for age determination, and relevant metadata such as length, weight, sex and maturity were registered.

### Sample handling and DNA extraction

Eight individuals (4 from Lofoten, 4 from Sørøya) were randomly selected for analysis of the intestinal content vs. mucosal layer. These intestinal samples were split open lengthwise, before the gut content was gently removed using a sterile disposable spatula, after which the mucosal layer was scraped off the gut lining with another spatula. For each of the remaining 14 individuals (8 from Lofoten, 6 from Sørøya), the gut content and mucosal layer were combined. For each extraction source (gut content, mucosa or mixture) per individual, three technical replicates were obtained for downstream processing (independent DNA extraction and PCR barcoding). Briefly, each individual sample was washed in 500 μl 100% EtOH and centrifuged before the ethanol was allowed to evaporate, after which dry weight was measured before proceeding to DNA extraction. Finally, the 22 individuals yielded eight gut content samples (in triplicate, G1-G3), eight mucosal samples (in triplicate, M1-M3) and 14 mixed samples (in triplicate, Mi1-Mi3). We also included one positive control consisting of DNA extract from an *Escherichia-Shigella* colony and two negative controls (extraction blanks) for a total of 93 samples. DNA was extracted from between 10 and 640 mg dry weight of gut content or mucosa using the *MoBio Powersoil HTP 96 Soil DNA Isolation Kit* (MoBio Laboratories, Carlsbad, CA, USA) according to the DNA extraction protocol (v. 4.13) utilized by the Earth Microbiome Project [26]. DNA was eluted in 100 μl Elution buffer, and stored at −20° Celsius.

### Sequence data generation

The V4 region of the 16S rDNA gene was amplified according to a slightly modified version of the EMP protocol (original v. 4.13) [26] using the primers 515f (5′-GTGCCAGCMGCCGCGGTAA-3′) and 806r (5′-GGACTACHVGGGTWTCTAAT-3′), the latter containing a 12 bp barcode unique for each sample (Additional file 2). Reactions were amplified for 30 cycles (3 min at 94°C, 30 cycles of 45s at 94°C, 60s at 52°C, and 90s at 72°C), with a primer extension of 10 min at 72°C. In contrast to the original EMP protocol, each extraction template was amplified *once*, including the negative extraction controls. PCR amplification was verified on a 1% agarose gel. The PCR products were normalized using the *SequalPrep Normalization Plate Kit* (Invitrogen, Paisley, UK) eluted in 20 μl Elution Buffer and the 96 Illumina compatible libraries were pooled. The pool was concentrated and purified using the *Amicon Ultra-0.5 mL Centrifugal Filter 30K Device* system (Merck KGaA, Darmstadt, Germany). Finally, the amplicons were paired-end sequenced by StarSEQ (StarSEQ GmbH, Mainz, DE) using Illumina MiSeq (Illumina, San Diego, CA, USA) V3 chemistry with the addition of 20% PhiX, allowing a maximum of 1 bp mismatch during demultiplexing. All sequence data have been deposited in the European Nucleotide Archive (ENA) under study accession number PRJEB22384 [27].

### 16S rRNA amplicon processing

Read qualities were assessed using *FastQC* [28] and primer and adapter sequences were removed using *cutadapt* [29] (--cut; hard cutoff of 48 first base pairs). Remaining phiX sequences were removed by mapping reads to the phiX reference genome [GenBank:J02482.1] using *BWA* [30] and discarding matching sequences using *seqtk* [31]. Downstream processing and sequence analysis was performed using *Mothur* (v. 1.36.1) [32] according to the Mothur Illumina MiSeq SOP. Briefly, paired-end reads were joined using the command *make.contigs* and filtered based upon a minimum average Phred quality score of 23, a maximum contig length of 375 bp and a zero-tolerance to ambiguous base pairs. Chimeras were removed using the *UCHIME* algorithm [33] implemented in *Mothur*, before the sequences (V4 region) were aligned and classified using the Silva SEED bacterial database (v. 119) as reference [34]. The dataset was clustered into OTUs based on a 97% sequence similarity using average neighbor clustering. The OTU table output of *Mothur* was imported into *Rstudio* (v. 0.99.893) [35] (based on R (v. 3.2.3) [36]) for further processing using the R package *phyloseq* (v. 1.14.0) [37]. Results were visualized using the R package *ggplot2* (2.0.0) [38].

OTU counts were normalized using a common scaling procedure, following McMurdie & Holmes 2014 [39]. This involves multiplying every OTU count in a given library with a factor corresponding to the ratio of the smallest library size in the dataset, replacing rarefying (i.e. random sub-sampling to the lowest number of reads). Normalizing using this procedure effectively results in the library scaling by averaging an infinite number of repeated sub-samplings. We excluded all OTUs not exceeding 0.001% of the total number of reads in the dataset in at least one sample. This corresponds to a minimum abundance threshold of 64 reads per OTU. Rarefaction curves, alpha diversity measures and ordination distances for non-metric multidimensional scaling (NMDS) plots were calculated in *phyloseq*. Distances were calculated based on OTUs with more than 10 reads (>99% of total number of reads).

### Oligotyping

Identification of putative bacterial strains and investigation of any concealed diversity was performed by oligotyping [40]. This is a supervised computational method to distinguish closely related but distinct bacterial organisms. In short, an oligotype is a concatenation of the most information-rich SNPs in a genetic marker sequence, and may be used as an approximate to a bacterial strain. A Shannon entropy analysis of the aligned 16S rRNA V4 sequences identifies information-rich positions, and a user-selected set of these are chosen for further analysis by visual inspection according to the Eren 2013 paper [40]. Oligotype profiles are then generated through a process involving different filtering parameters. Due to the supervised nature of the method, we limited the oligotyping to the ten most highly abundant OTUs in our dataset. The following parameters were applied: s = 4, a = 0.1, A = 0 and M = 176 (mean library size/1000).

### Statistical analysis

Differences in within-individual diversity (alpha diversity) were studied using a linear mixed effects model with random effect for fish, following Zuur et al. 2009, chapter 5.3 [41]. The “optimal” models (i.e. the models that best describe the individual diversities) were identified by a top-down reduction strategy and selected through t-tests based on Restricted Maximum Likelihood (REML) estimation. Model assumptions were verified through plotting of residuals. The effect size of the variables on the diversity measures was determined by comparing the estimate of that variable to the interquartile range (middle 50% of the spread) of the diversity measure. For the sake of simplicity, we defined the statistical variable “tissue” to contain both mucosal tissue and gut content, although the latter in not a tissue per se. Differences in bacterial community composition (beta diversity) between gut content and mucosal tissue, and between the two locations, were assessed using Permutational Multivariate Analysis of Variance (PERMANOVA) using the *adonis* function in the R package *vegan* [42]. The test was run with 20,000 permutations, applying both OTU-based (Bray-Curtis/Jaccard) and phylogeny-based (weighted and unweighted UniFrac) distance measures. For location analysis, the mean normalized abundance for each of the 139 OTUs in the 22 individuals was applied. For tissue analysis, normalized abundance of all gut and mucosa replicates (n = 42) was used. Due to dependency within replicates for each fish, individuals were used as strata (blocks) in the gut vs. mucosa testing. PERMANOVA assumes the multivariate dispersion in the compared groups to be homogeneous; this was verified using the *betadisper* function (*vegan*). Similarity percentage (SIMPER) procedure implemented in *vegan* was used to quantify the contribution of individual OTUs to the overall Bray-Curtis dissimilarity between gut content and mucosa. OTUs with a statistically significant differential abundance between the locations were determined using *MetaStats* [43]. Only OTUs containing more than 0.1% of the total number of reads in the normalized and filtered dataset (of 139 OTUs) were used in this analysis. Finally, differences in relative abundance of the individual oligotypes between Lofoten and Sørøya were studied using linear mixed effects models for log-transformed read counts, and selected and verified as described above, with individuals and oligotypes (within individuals) treated as random effects.

## Results

We obtained 14,944,840 paired-end reads that were assembled based on sequence overlap. Of the original 93 samples, this dataset contained 86 cod and three control libraries (four replicates failed to sequence). The libraries (excl. controls) had a median size of 173,377 reads per replicate (Additional file 3). The positive control contained 96.5% sequences classified as *Escherichia-Shigella*, as expected. The negative controls yielded less than 0.01% of total number of reads in the dataset (Additional file 4). These reads represent known reagent and laboratory contaminant OTUs [44]. Due to insufficient sequencing of a Sørøya sample (S04, all three replicates), only seven individuals (4 Lofoten, 3 Sørøya) were used for comparison of diversity in gut and mucosa.

After normalizing OTU abundances to 74,374 reads, and filtering of OTUs based on abundance, 139 OTUs representing 98.6% of the dataset were identified (Additional file 5). Rarefaction curve analyses on the normalized data show that variation in the number of OTUs detected per sample is not caused by uneven sequencing depth (Fig. 1b). The technical replicates cluster close to one another per respective tissue type and individual specimen in a multivariate NMDS analyses (Fig. 1c), and this methodological consistency is reflected by the similar OTU composition of the individual replicates (Additional file 6). Finally, we observe a large overlap between the OTU community profiles when clustering individuals from both locations together (Fig. 1d).

### Taxonomic composition of the bacterial gut community

The 139 OTUs are binned into 9 phyla, 14 classes and 20 orders (Fig. 2, Additional file 7), with all the abundant OTUs having been identified at both locations. 23 OTUs are detected in one of the locations only (Fig. 2), of which 17 belong to the 30% least abundant OTUs (Additional file 5). A core microbiome of 23 OTUs is identified based on shared membership in all 22 individuals (Fig. 2, Additional file 8). This core community includes the ten most abundant OTUs in the dataset, and represents about 93% of the total number of reads. The *Proteobacteria* represents 67% of the total read count (63 OTUs), followed by *Fusobacteria* (15.7% / 5 OTUs), *Firmicutes* (6.1% / 33 OTUs) and *Bacteroidetes* (4.5% / 18 OTUs). *Gammaproteobacteria* represents the major bacterial class, with a median relative abundance of 64.6% among the samples. This is mainly due to the *Vibrionales* order; with a relative abundance ranging from 27% - 97% it is clearly the most prolific member of the intestinal microbiome in northern coastal Atlantic cod (Fig. 3a). *Fusobacteriales*, *Clostridiales* and *Bacteroidales* are abundant in some, but not all individuals.

**Figure 2:**
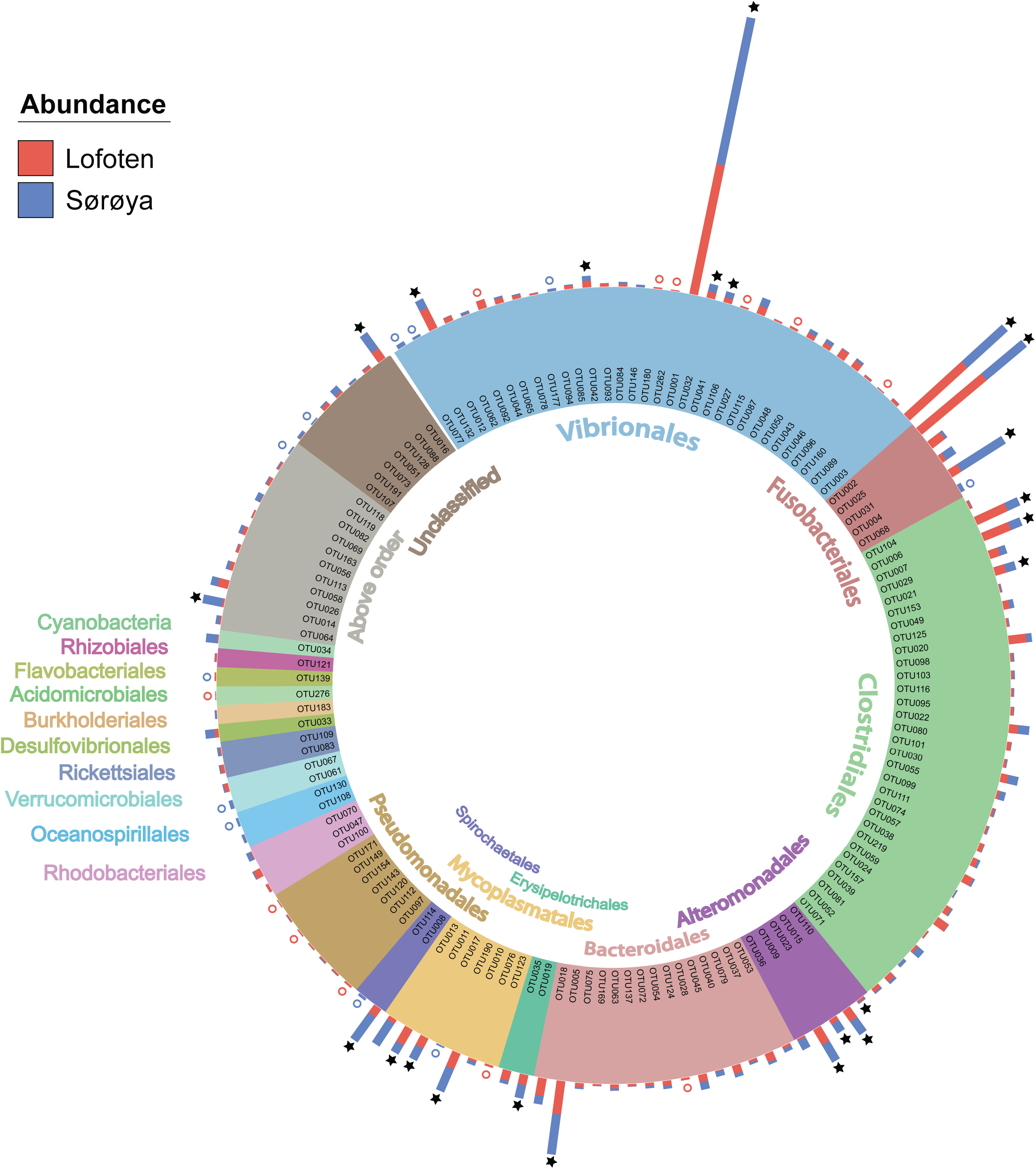
Classification of the Atlantic cod intestinal bacterial community members and their relative abundances at two sample locations. The 139 OTUs are grouped based on an order-level classification (color). Stacked bars represent read abundance (square root transformed) of each OTU in Lofoten (red) and Sørøya (blue). Some OTUs are found in all specimens (stars), whereas others (colored circles) are observed in only one of the locations. OTU order classifications are given in the inner and outer part of the circle.

**Figure 3:**
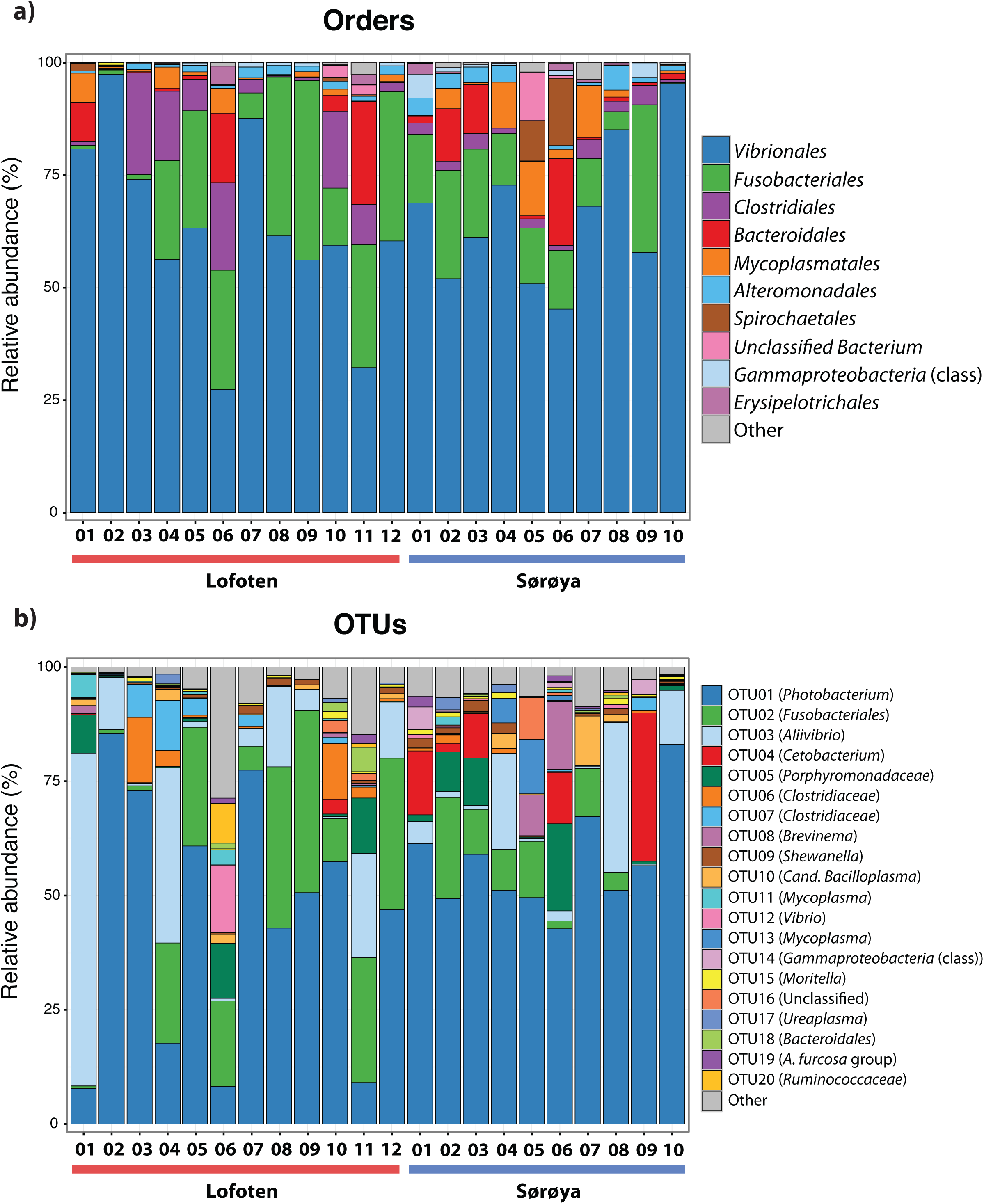
Taxonomic composition of the intestinal microbiome in Atlantic cod specimens from Lofoten and Sørøya. **a)** Relative abundance of reads classified to bacterial orders. Colors represent the 10 most abundant orders **b)** Relative abundance of reads by OTUs (at 97% similarity) classified to lowest identifiable taxonomic level. Remaining orders or OTUs are merged into “Other” categories (grey bars). The relative abundances of replicate samples or tissues were averaged per individual.

### Dominant members of the gut microbiome

The ten most abundant OTUs in the Atlantic cod individuals represent 88.6% of the total number of sequence reads (Table 1). The most abundant OTU (OTU 01), a *Photobacterium*, represents 78% of all *Vibrionales* reads, and comprises 50% of all sequences in the study. It also constitutes > 50% of the reads in 13 of the 22 specimens (Fig. 3b). In five of the seven specimens used for analysis of differences between gut content and mucosa, OTU 01 represents > 45% of the reads (mean of triplicates) in both gut content and mucosa (Additional file 9). Blast results provide no clear species-level identification of this OTU due to multiple entries of different species. In the cases that OTU 01 is not the most prolific member, other common OTUs (such as OTU 03 (*Aliivibrio*), or OTU 02 (*Fusobacteriales*)) show a higher abundance (Fig. 3b).

**Table 1:** The 10 most abundant OTUs in the Atlantic cod samples. The table shows the OTU name, the lowest possible classified taxonomic level, the number of reads after normalization and the relative abundance of the OTU in the cod community.

| OTU nr. | Taxonomy | Total reads | Relative abundance (%) |
| --- | --- | --- | --- |
| 01 | *Photobacterium* | 2 436 281 | 50.36 |
| 02 | *Fusobacteriales* | 635 730 | 13.14 |
| 03 | *Aliivibrio* | 576 357 | 11.91 |
| 05 | *Porphyromonadaceae* | 161 547 | 3.46 |
| 04 | *Cetobacterium* | 167 536 | 3.34 |
| 06 | *Clostridiaceae* | 85 810 | 1.77 |
| 07 | *Clostridiaceae* | 65 527 | 1.35 |
| 08 | *Brevinema* | 59 540 | 1.23 |
| 10 | *Cand. Bacilloplasma* | 43 301 | 1.12 |
| 09 | *Shewanella* | 54 358 | 0.89 |

### Intestinal microbial diversity

The individual tissue samples contain between 34 - 90 OTUs (Fig. 4), and vary also in diversity estimated by Shannon (H) and Inverse Simpson (1/D) indexes (Additional files 10 and 11). The optimal linear mixed effects models for the three diversity measures, including fish specimen as a random effect, reveal a statistically significant difference between mucosa and gut content (Table 2, Additional file 12). For all three measures, the diversity is significantly lower in mucosa than in the gut. However, the estimated differences are small compared to the variation in the diversity measures between all 22 individuals. For the number of OTUs, the estimate in Table 2 corresponds to 23% of the interquartile range for the variation in OTUs between all fishes, while for the Shannon and Inverse Simpson indexes the estimates correspond to 26% and 42%, respectively, of the interquartile range. The reason why such small differences become significant, is that we have triplicates of both gut and mucosa samples for seven individuals (Additional file 9), ensuring high statistical power for the comparison of gut vs. mucosa. For the individuals with mixed tissue triplicates, we do not have any other types of tissue samples. Therefore, we have much lower statistical power for comparing mixed tissue with the other tissue types. As a result, we do not find a significant effect of mixed tissue even though the estimates for two of the diversity measures are of the same order of magnitude as for mucosa. The results in Table 2 also show that length and sex of the Atlantic cod specimens have a significant impact on the observed number of OTUs. There are no significant differences in within-sample diversity between the two locations (Additional file 12).

**Figure 4:**
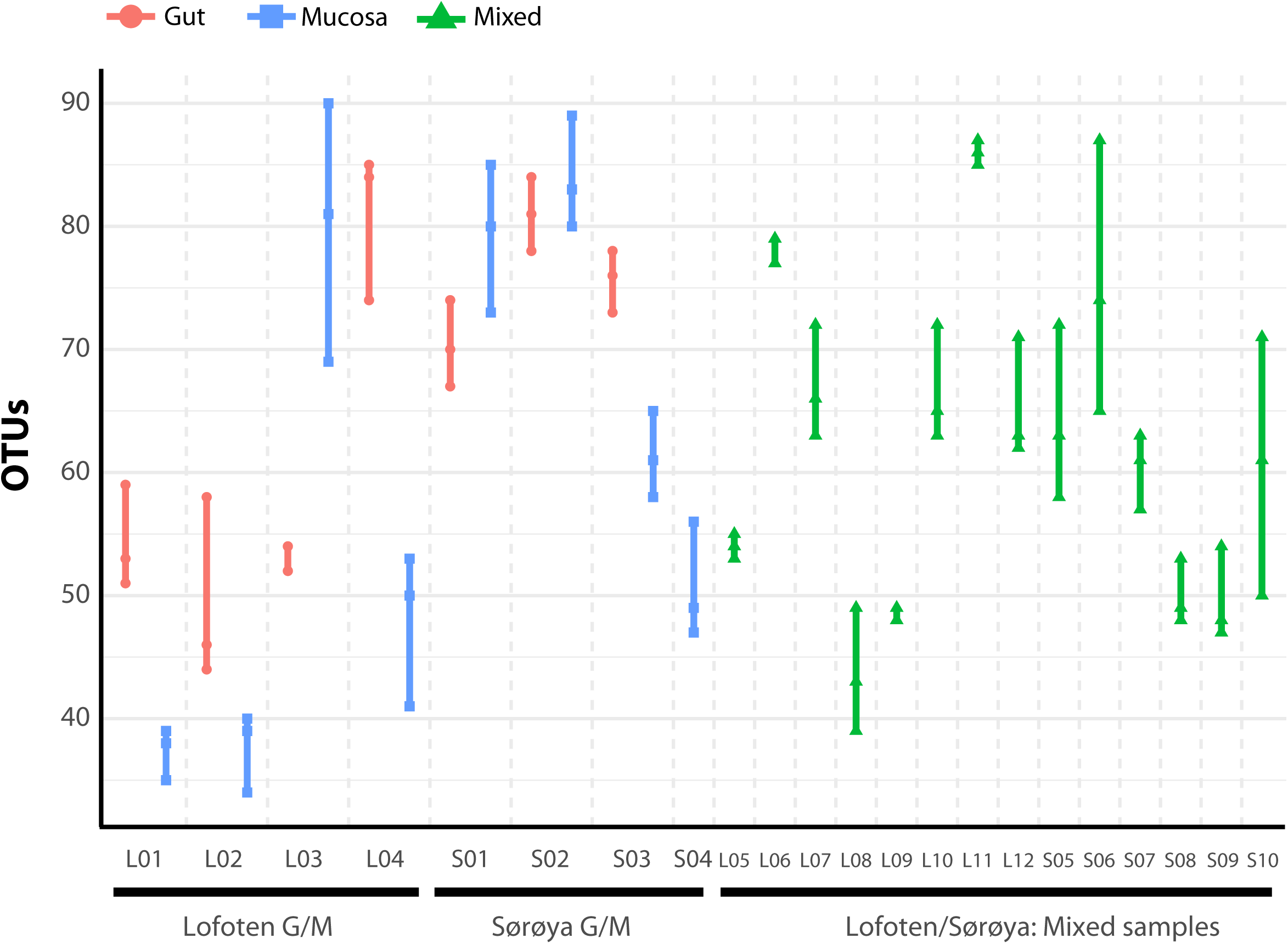
Number of observed OTUs in 22 Atlantic cod samples. Each of the 86 replicates is represented by a point. Individuals sampled for both gut (red) and mucosa (blue) on the left half of the figure, individuals sampled for a mix (green) of gut and mucosa on the right half.

**Table 2:**
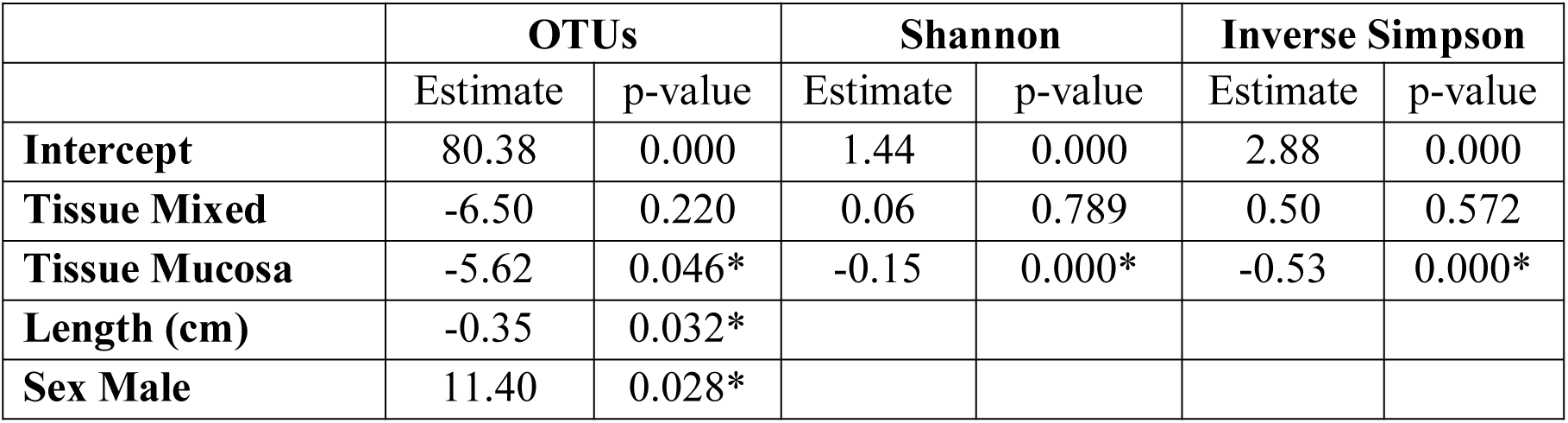
Effects of covariates on intestinal diversity (alpha diversity). Results from the optimal linear mixed effects models used to analyze within-sample (alpha) diversity. “Estimate” indicates the estimates of the regression coefficients for the fixed effects in the model. Tissue estimates are given relative to gut, and the sex estimate is given relative to females. Significant effects (p < 0.05) of covariates are indicated by an asterisk.

Results from between-sample (beta) diversity analyses agree with those from the within-sample (alpha) analyses. PERMANOVA analysis reveals a statistically significant difference in community composition between gut and mucosa using Bray-Curtis, Jaccard and Weighted UniFrac diversity measures, while there is no significant difference between Lofoten and Sørøya (Table 3). Nevertheless, the difference between gut content and mucosa has a very small effect size, and can only explain about 1 - 3% of the total variation in the gut and mucosal samples. The reason why these small effects become significant is, as described above, that we have high statistical power for comparison of gut vs. mucosa. The significant difference between gut content and mucosa is preserved when limiting the analysis to the five most abundant OTUs (Additional file 13), indicating that a restricted number of highly abundant OTUs is responsible for the differences. This is in agreement with results from the SIMPER analysis, where the five most abundant OTUs (OTU 01-05) together contribute 77.3% to the observed (Bray-Curtis) dissimilarity between gut and mucosa (Additional file 14). MetaStats analysis of differential abundance between the locations identifies four OTUs – *Fusobacteriales* (OTU 02), *Cetobacterium* (OTU 04), *Gammaproteobacteria* (OTU 14) and *Shewanella* (OTU 23)-with a low p-value (Table 4), although no significance remains after Bonferroni correction.

**Table 3:** PERMANOVA analysis of diversity differences between bacterial communities (beta diversity). Location: 21 degrees of freedom (df), tissue: 41 df. Significant effects (Bonferroni corrected, p < 0.00625) are indicated by an asterisk.

|  |  | Bray-Curtis | Jaccard | Unweighted UniFrac | Weighted UniFrac |
| --- | --- | --- | --- | --- | --- |
| **Location** | R2 | 0.11 | 0.09 | 0.07 | 0.07 |
|  | p-value | 0.047 | 0.041 | 0.146 | 0.218 |
| **Tissue** | R2 | 0,01 | 0.01 | 0.03 | 0.01 |
|  | p-value | 0.001* | 0.001* | 0.015 | 0.003* |

**Table 4:**
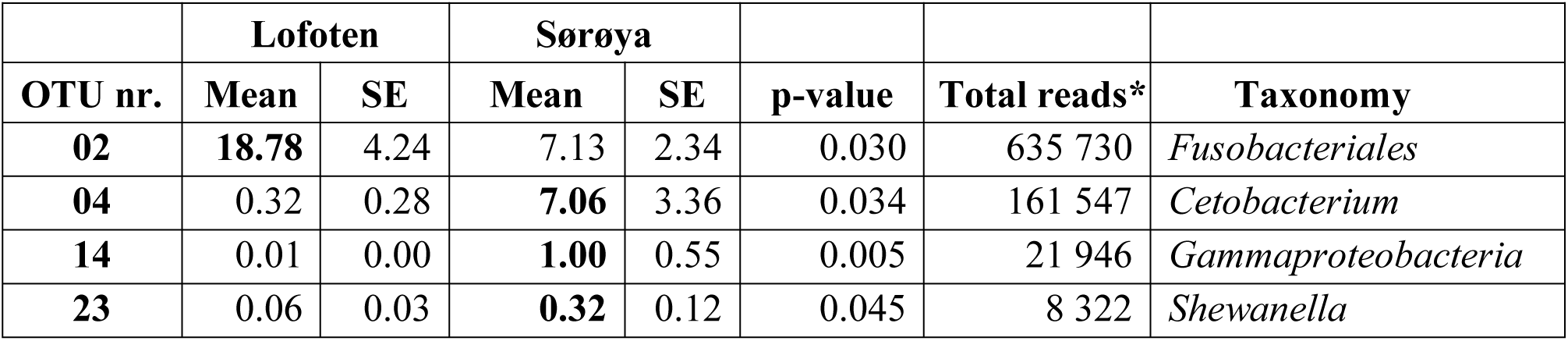
OTUs differentially abundant between Lofoten and Sørøya. 41 OTUs, each representing more than 0.1% of the total abundance, were analyzed using *MetaStats*. Mean and SE values are in percent. None of the OTUs give p-values below the Bonferroni corrected p-value (0.05/41 = 0,001) significance threshold. Bold values mark the location of highest mean abundance. *Total abundance of OTU in all samples.

|  | Lofoten |  | Sørøya |  |  |  |  |
| --- | --- | --- | --- | --- | --- | --- | --- |
| OTU nr. | Mean | SE | Mean | SE | p-value | Total reads* | Taxonomy |
| **02** | **18.78** | 4.24 | 7.13 | 2.34 | 0.030 | 635 730 | *Fusobacteriales* |
| **04** | 0.32 | 0.28 | **7.06** | 3.36 | 0.034 | 161 547 | *Cetobacterium* |
| **14** | 0.01 | 0.00 | **1.00** | 0.55 | 0.005 | 21 946 | *Gammaproteobacteria* |
| **23** | 0.06 | 0.03 | **0.32** | 0.12 | 0.045 | 8 322 | *Shewanella* |

### Identification of oligotypes with differential abundance

Oligotyping reveals higher taxonomical detail in six of the ten most abundant OTUs (Fig. 5a). Each of the OTUs 02, 04 and 06 - 09 consist of at least two oligotypes, while the remaining four OTUs show no such substructures (i.e. there is one oligotype per OTU). Of the six OTUs with multiple oligotypes, OTU 04 and OTU 09 contain oligotypes with a differential abundance between the two locations. OTU 04 oligotype “C” has a small but significantly higher abundance in Sørøya than in Lofoten (Table 5). P-values just above 5% of an additional OTU 04 oligotype (“T”) and two OTU 09 oligotypes (“ATGAG/CGAGT”) weakly indicate a differential abundance between the two locations. The remaining OTUs with multiple oligotypes contain no differentially abundant oligotypes (Fig. 5b).

**Figure 5:**
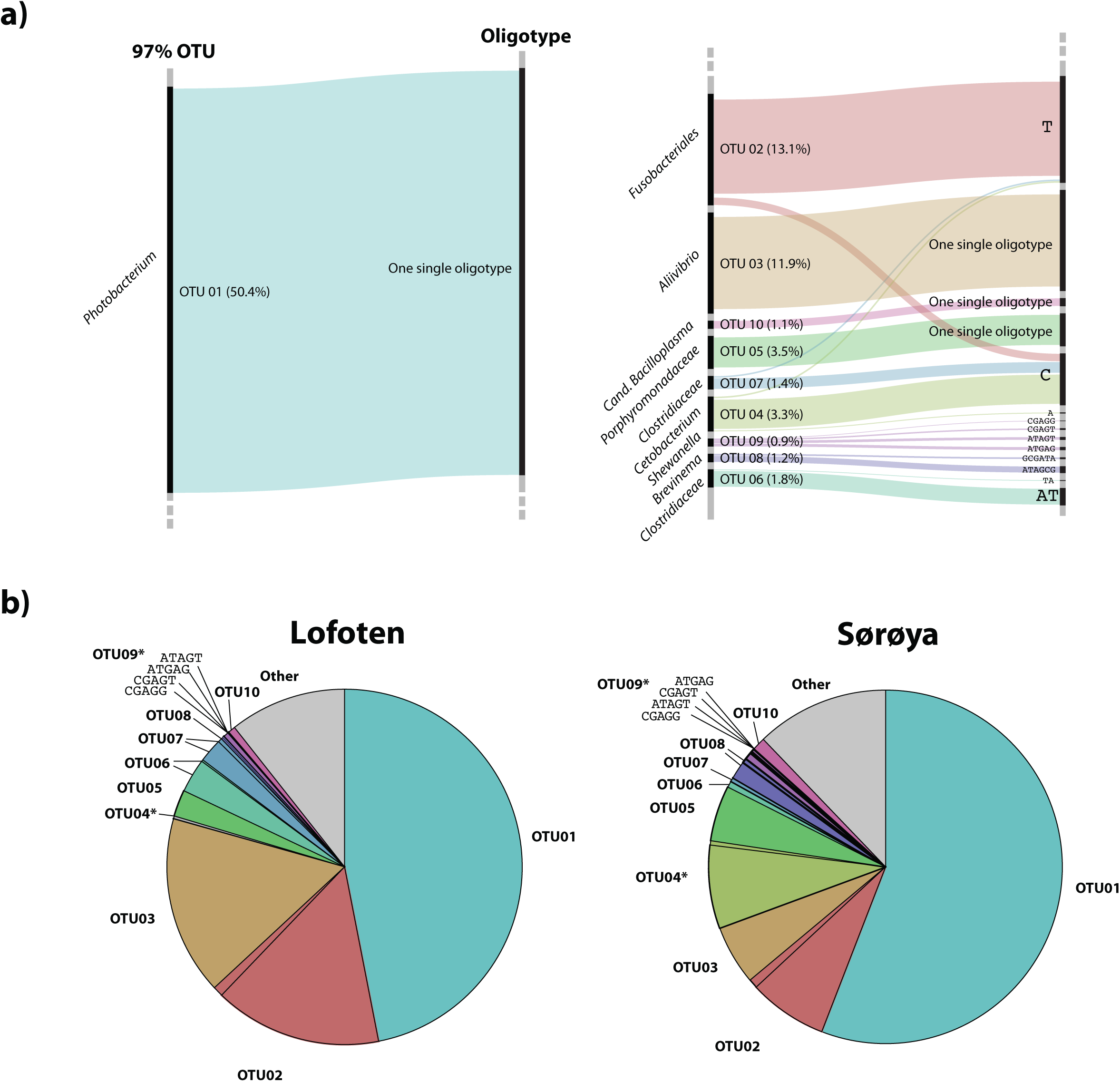
Oligotypes among the most abundant OTUs. **a)** Substructure of the ten most abundant OTUs in the gut microbiome of Atlantic cod. Black bars on left side represents OTUs, while bars on the right side represents unique oligotypes. The height of a colored bar represents its overall abundance, also stated in the parenthesis behind each OTU. Genus (or lowest classified taxonomic level) is given to the left of each OTU. For four OTUs, no additional oligotypes are detected, and the most abundant OTU (OTU01, left) consists of a single oligotype. **b)** Relative abundance of OTUs and their representative oligotype(s) in the gut microbiome of Atlantic cod in two locations. Colors represent different OTUs, and identically colored sectors represent different oligotypes within an OTU. OTUs containing differentially abundant oligotypes as determined by mixed effects modeling are marked with an asterisk (*). For illustrational purposes, the four oligotypes of OTU09 are labelled in the figures.

**Table 5:** Differential abundance of individual oligotypes. “Estimate” indicates the estimated effect of Sørøya compared Lofoten. The estimates are obtained from a linear mixed effects models for log-transformed read counts (with one added to all read counts) with individuals and oligotypes (within individuals) as random effects, and tissue type, oligotype, location and interaction between oligotype and location as fixed effects. OTU04 oligotype C has a small but significantly higher abundance in Sørøya than in Lofoten (p < 0.05, *). P-values just above 5% of three other oligotypes (.) weakly indicate a differential abundance between Lofoten and Sørøya.

| OTU nr. | Genus | Oligotype | Estimate | p |
| --- | --- | --- | --- | --- |
| 04 | *Cetobacterium* | C | 3.37 | 0.003* |
|  |  | T | 1.70 | 0.051 . |
|  |  | A | -0.84 | 0.316 |
| 09 | *Shewanella* | ATGAG | 1.77 | 0.055 . |
|  |  | ATAGT | -1.00 | 0.309 |
|  |  | CGAGT | 1.70 | 0.057 . |
|  |  | CGAGG | 0.44 | 0.603 |

## Discussion

Here, we have investigated the diversity and taxonomic composition of the gut microbiome of Atlantic cod from two separate locations along the Norwegian coastline based on 16S rRNA sequencing. The Atlantic cod gut community is dominated by *Proteobacteria*, *Fusobacteria*, *Firmicutes* and *Bacteroidetes* (Additional file 7); all known to be among the most dominant phyla in the gastrointestinal tract of marine fish [5,12]. *Vibrionales*, *Fusobacteriales*, *Clostridiales* and *Bacteroidales* are among the most abundant orders. In particular, the order *Vibrionales* occurs in high abundance (64% of reads), which is in agreement with what was found in a meta-analysis of marine fish microbiomes [4] and in the intestinal microbiome of 11 wild Atlantic cod individuals caught in a single location in the Oslo fjord [24]. While this study reported large inter-individual differences in OTU abundance [24], these differences were largely drive by variation in abundance of the rare members of the microbial community. Yet – similar to what we report here – the Oslo fjord individuals shared a low number of highly abundant OTUs, with *Vibrionales* representing > 50% of the total number of reads [24]. Unlike previous findings, however, we identify a single *Vibrionales* 16S rRNA oligotype (genus *Photobacterium*, OTU 01) that comprises 50% of the sequencing data. To our knowledge, the occurrence of a single, prolific oligotype over such a large spatial scale has not been reported before in marine fish.

There are two alternative explanations why we observe no genetic diversity within this abundant oligotype: It is possible that the 16S rRNA V4 region does not capture sufficient sequence variation to resolve fine-scaled population structure within the *Vibrionales* that are part of the Atlantic cod gut microbiome. Alternatively, the presence of a single, dominant *Vibrionales* oligotype represents the colonization and subsequent numerical increase of a single -or closely related- bacterial strain. While at this stage it is difficult to confidently exclude the first explanation, there are several observations that indicate that this ubiquitous strain abundance may reflect a biological reality rather than a methodological limitation. First, we find another highly abundant *Vibrionales* OTU (OTU 03) that also has a lack of within OTU diversity (Fig. 5a). This indicates that the V4 region is capable of distinguishing various *Vibrionales* strains down to the genus level. Second, other studies have also observed the occurrence of a low number of highly abundant taxa (including *Photobacterium*) in the fish intestinal microbiome, although these did not resolve taxonomy down the level of oligotypes. For example, the gut microflora of three shark species is dominated by a single *Photobacterium* OTU that represents up to 98% of the total shared sequences [45]. Similarly, 93% of the data comprised a single *Photobacterium* species in the intestines of two wild cold water adapted notothenioid species of the Antarctic Ocean [46]. Moreover, four captive adults of Atlantic halibut were shown to contain > 70% of luminous *Photobacterium phosphoreum* isolates through culture-dependent methods [47]. Finally, little differentiation was observed in the Atlantic salmon gut microbiome from populations on both sides of the Atlantic Ocean [13]. This indicates that specific bacterial strains – in particular *Photobacterium* – may be efficient dispersers over large spatial scales [48,49]. These observations suggest that *Photobacterium* is especially capable of marine dispersal and subsequent colonization and proliferation in the Atlantic cod gut. The latter is supported by the fact that the *Photobacterium* OTU 01 is found in high relative abundance (> 45%) in the mucosa in five of the seven Atlantic cod samples used for analysis of gut content vs. mucosa.

We observe a small, but significant difference between the microbial communities in gut content and mucosal tissue. The alpha diversity estimates reveal a slightly less diverse community in mucosa than in gut content, which is in agreement with previous findings in fish [22,50]. This observation supports a hypothesis assuming that the intestinal mucosa hosts a subset of specialized bacteria compared to what is present in the more heterogeneous gut content of an omnivore. We find that the numerically dominant OTUs are abundant in both gut and mucosa. From this, we derive that these OTUs are associated with the intestinal mucosa, as part of the residential (autochthonous) microbiota. This permanent community is most likely to interact with the host and may serve functionally important purposes, i.e. protective immunity [2]. Nevertheless, the limited community differences we observe between the tissue types may also reflect our sampling methodology; out of necessity, the samples were stored in ethanol for a prolonged amount of time, and separation of mucosa and gut content would most likely have been more efficient by sampling of fresh tissue.

Based on OTU level classification, we find no significant overall bacterial community differentiation between Atlantic cod from Lofoten and Sørøya. The intestinal microbiome in coastal Atlantic cod from Lofoten and Sørøya may be similar due to several reasons. First, a strong host-microbe interdependence based on mutual benefits between Atlantic cod and specialized, residential bacteria would promote congruent gut communities in both locations. This requires Atlantic cod to actively recruit certain bacterial species; such host selection is suggested in adult zebrafish and Atlantic salmon parr [51,52]. Second, environmental factors such as diet, temperature or water bacterioplankton content common for the two locations may contribute to a similar diversity and community structure of the fish gut microbiomes (reviewed in [5,53]). While no OTU level differentiation in beta diversity is detected between these two locations, two OTUs contain oligotypes that show limited spatial differentiation that could reflect the limited acquisition of more “local” strains into the Atlantic cod microbiome. Hence, oligotyping may reveal a level of spatial segregation which a prevailing method could not have detected [40].

The Atlantic cod gut microbiome contains considerable amounts of *Fusobacteriales*, *Cetobacterium, Aliivibrio*, *Porphyromonadaceae*, *Clostridiaceae*, *Brevinema* and *Shewanella*, which have been observed in fish intestines in other studies [54–57]. However, *Photobacterium* (OTU 01) is the predominant oligotype in both gut and mucosal samples from both locations. A high abundance of this oligotype in mucosa suggests it is a residential bacterium associated with the gut lining, and thus potentially involved in host-microbial co-evolution. For example, some *Photobacterium* species show antagonistic activity towards bacterial pathogens in Atlantic cod, *Vibrio anguillarum* and *Aeromonas salmonicida* [58] and hence contribute to protective immunity in Atlantic cod. Considering the loss of the MHC II pathway of the adaptive immune system, the Atlantic cod could therefore benefit from housing *Photobacterium* in its intestines.

Atlantic cod is a dietary generalist that consumes a varied diet. Such dietary behavior might intuitively lead to an expectation of an associated diverse intestinal microbiota, given an exposure to more bacterial diversity. Our lack of large-scale population differentiation suggests this is not the case in Atlantic cod. Interestingly, a negative association between diet diversity and microbial diversity has been observed in stickleback and perch [59]. It was hypothesized that generalists have more nutritionally diverse gut environments that sustain a limited number of competitively dominant bacteria at high abundance. Those individuals exposed to a more varied diet are shown to have a substantial increase of *Gammaproteobacteria*. This bacterial class also occurs in high relative abundance in our Atlantic cod samples (i.e. *Photobacterium*) and hence this observed microbiome may indeed reflect that of a dietary generalist.

Another explanation for the lack of differentiation in the microbiome of the Atlantic cod intestine may be related to its lack of MHC class II, a key pathway in the vertebrate adaptive immune system. MHC II is produced in antigen-presenting cells which phagocytize extracellular particles, including bacteria. It is therefore proposed that this pathway is involved in the recognition and management of a complex bacterial community [60]. The absence of such a regulatory mechanism may lead to a limit in the number of resident bacterial species that can be maintained in its intestines. Indeed, it has been suggested that a lower microbial diversity observed in invertebrates is due to their lack of an adaptive immune system [61]. Thus, the role of the Atlantic cod immune system in the active maintenance of microbial species requires further investigation.

## Conclusions

In this study, we find that the Atlantic cod intestinal microbiome is dominated by a limited number of abundant 16S rRNA oligotypes, with limited differentiation between intestinal bacterial communities in Lofoten and Sørøya. A significant yet small difference in the community diversity between gut and mucosa suggests that the abundant members of the microbiome are part of the more permanent inhabitants of the gut, that possibly play a role in host-microbe co-evolution. Finally, a single *Photobacterium* oligotype is particularly abundant in the Atlantic cod gut microbiome in the two locations 470 km apart. Future studies involving metagenomic/-transcriptomic shotgun sequencing will provide more detailed insights into the diversity and functional capacity of the intestinal microbiome of Atlantic cod.

## Availability of supporting data

The data set supporting the results of this article is available in the European Nucleotide Archive (ENA), study accession number PRJEB22384, http://www.ebi.ac.uk/ena/data/view/PRJEB22384 [27].

## Authors’ contributions

SJ, BS and THA conceived and designed the experiments. KSJ provided the initial framework for the study. ESR and SJ sampled the specimens. ESR performed the laboratory work. ESR and THA performed data analysis. ØB, THA, ESR and BS interpreted the results. ESR and BS wrote the paper with input of all authors. All authors have read and approved the final manuscript.

## Acknowledgements

This work was funded by a grant from the Research Council of Norway (project no. 222378) and University of Oslo (Strategic Research Initiative) – both to KSJ. We thank Ole Christian Lingjærde for fruitful discussions. We also thank Børge Iversen and Helle Tessand Baalsrud for their kind help in sampling cod specimens in Lofoten, and Martin Malmstrøm, Paul Ragnar Berg and Monica Hongrø Solbakken for sampling in Sørøya.

## Additional files

### Additional file 1.xlsx: Metadata

Data collected for all cod specimens used in the study.

### Additional file 2.xlsx: Primers and barcodes

Primer sequences and unique barcode for each sample.

### Additional file 3.xlsx: Library sizes

Number of forward and reverse sequences per library, including raw reads, reads after removal of PhiX sequences and reads after the *Mothur* pipeline (used for downstream analysis).

### Additional file 4.xlsx: Positive and negative controls

Taxonomic composition and number of raw reads in the positive and the two extraction (negative) controls.

### Additional file 5.xlsx: OTU table and classification

Left part of table: Key information of each OTU after abundance filtering and common scaling, including total number of reads, overall relative abundance and the number of individuals each OTU occur in (frequency). Color codes are given below the table. Middle part: The final OTU table after abundance filtering, common scaling and merging of six replicates per sample into separate individuals. Right part: Classification of each OTU.

### Additional file 6.pdf: Relative abundance of OTUs in all 86 replicates

The 20 most abundant OTUs in cod are colored, while remaining OTUs are merged into the “Other” category (grey bars). G = Gut, M = Mucosa, Mi = Mixed.

### Additional file 7.xlsx: Taxa level information

Key relative abundance values of the different taxa (at all taxonomic levels) in individuals from Lofoten and Sørøya, and in the overall dataset.

### Additional file 8.xlsx: Core community

Core OTUs found in all individuals from Lofoten and Sørøya. Genus or lowest classified taxonomical level shown in the last column. c_ = class, o_ = order, f_ = family.

### Additional file 9.pdf: Relative abundance of OTUs in the gut (G) and mucosal (M) samples

All replicates from four Lofoten and three Sørøya individuals are shown. The 20 most abundant OTUs in cod are colored, while remaining OTUs are merged into the “Other” category (grey bars).

### Additional file 10.pdf: Shannon (a) and Inverse Simpson (b) diversity in the 22 cod samples

Each of the 86 replicates is represented by a point. Individuals sampled for both gut (red) and mucosa (blue) on the left half of the figure, individuals sampled for a mix (green) of gut and mucosa on the right half.

### Additional file 11.xlsx: Alpha diversity values

Alpha diversity estimates of the Atlantic cod intestinal microbial samples. All samples were normalized by a common scaling procedure (see Methods). OTUs were clustered based on 97% sequence similarity.

### Additional file 12.xlsx: Alpha diversity differences - Random intercept model

Results from the *beyond optimal* model and the final random intercept model fitted through model selection according to Zuur et al. 2009 chapter 5.7*, used in testing for significant differences in alpha diversities within locations and tissues. Significant values (< 0.05) are marked in bold.

*Zuur, A. et al., 2009. *Mixed Effects Models and Extensions in Ecology with R*, Springer. Available at: http://link.springer.com/book/10.1007%2F978-0-387-87458-6#section=15123&page=1.

### Additional file 13.xlsx: PERMANOVA with the five most abundant OTUs only

Results from PERMANOVA analyses of bacterial community differences using the five most abundant OTUs only. Bacterial community differences (beta diversity) were tested between the different locations, and between gut content, intestinal mucosa and a mixture of the two. P-values < 0.05 are marked in bold. These are still above the Bonferroni threshold (p = 0.00625).

### Additional file 14.xlsx: SIMPER results

SIMPER analysis results showing the contribution of each OTU to the Bray-Curtis dissimilarity between gut content and mucosa. In agreement with the results shown in Additional file 13, the five most abundant OTUs (OTU01 - 05) contribute 77.3% to the observed dissimilarity (see column “Cumulative contribution”). Significance information is supplied in the two last columns.

---

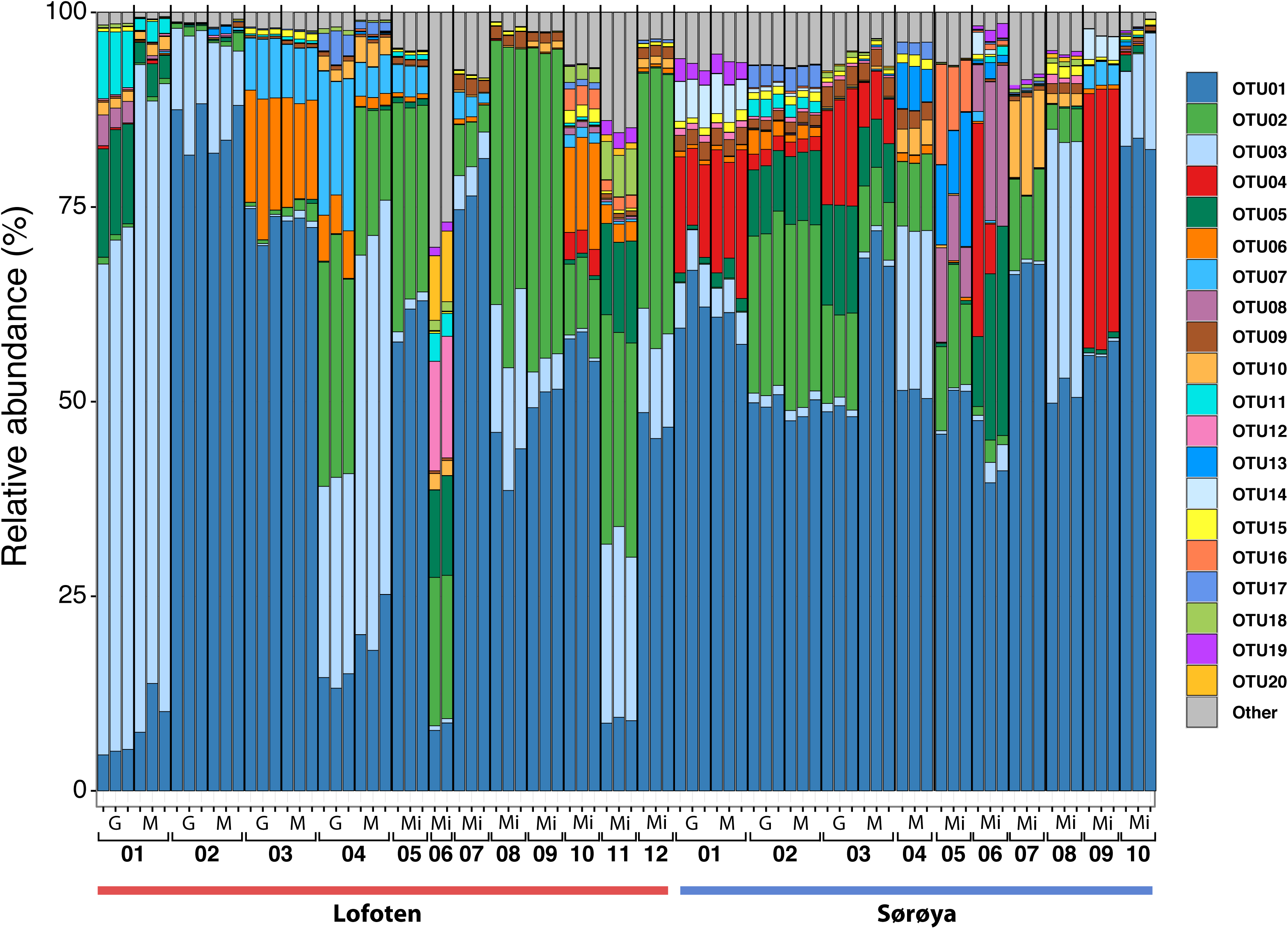

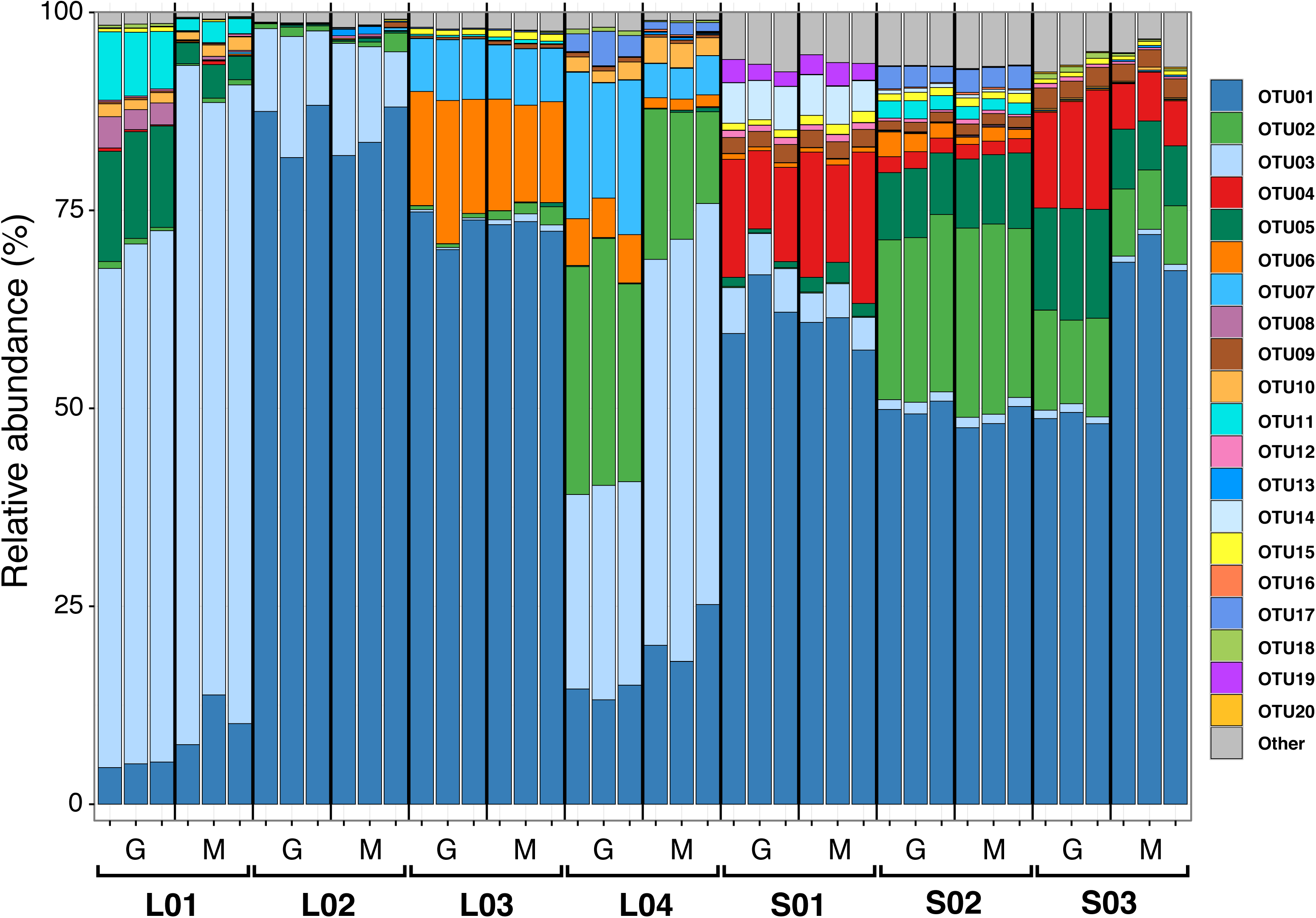

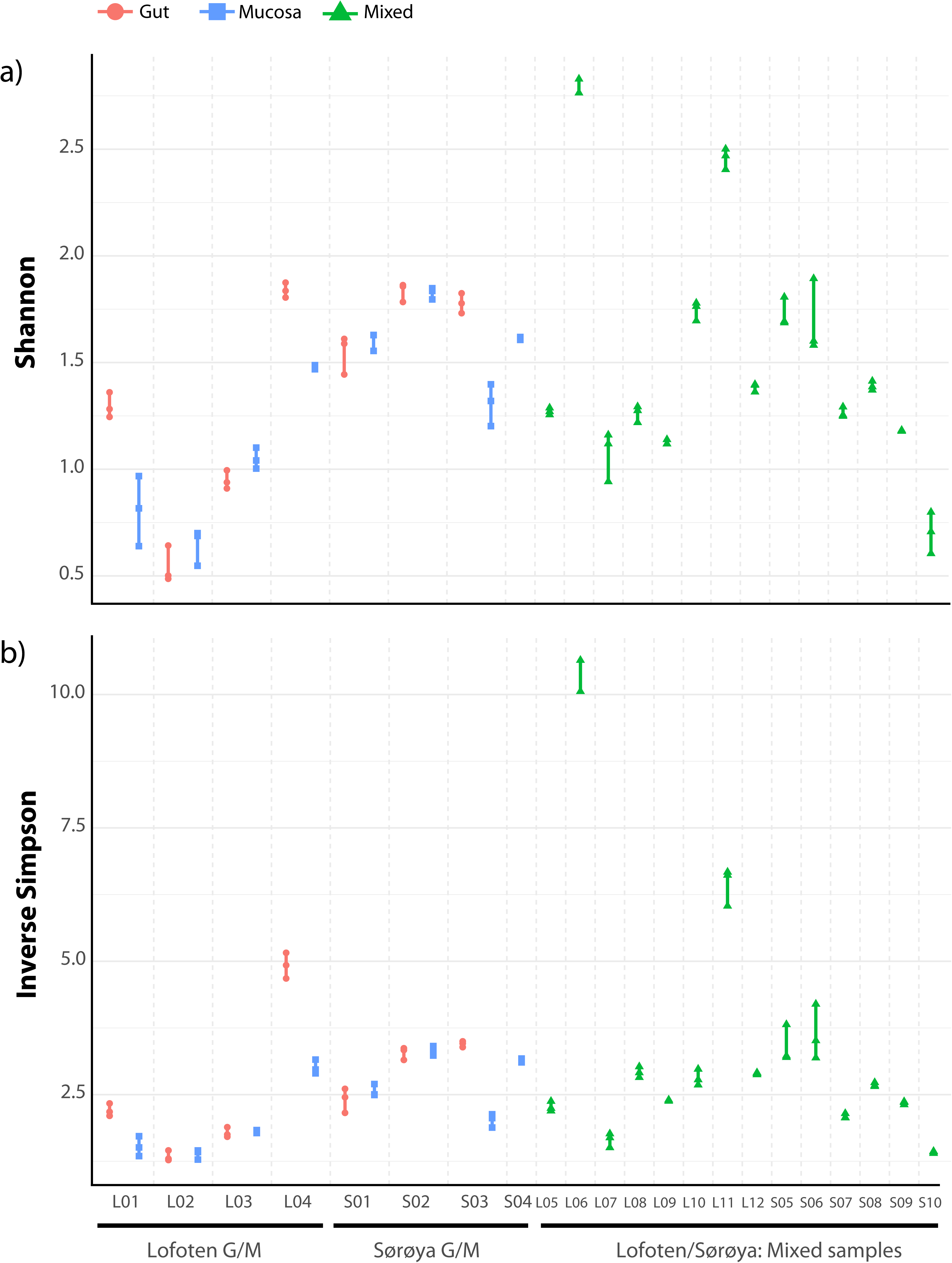

